# Smartphone-based Sickle Cell Disease Detection and Monitoring for Point-of-Care Settings

**DOI:** 10.1101/2020.05.11.087593

**Authors:** Shazia Ilyas, Mazhar Sher, E Du, Waseem Asghar

## Abstract

Sickle cell disease (SCD) is a worldwide hematological disorder causing painful episodes, anemia, organ damage, stroke, and even deaths. It is more common in sub-Saharan Africa and other resource-limited countries. Conventional laboratory-based diagnostic methods for SCD are time-consuming, complex, and cannot be performed at point-of-care (POC) and home settings. Optical microscope-based classification and counting demands a significant amount of time, extensive setup, and cost along with the skilled human labor to distinguish the normal red blood cells (RBCs) from sickled cells. There is an unmet need to develop a POC and home-based test to diagnose and monitor SCD and reduce mortality in resource-limited settings. An early-stage and timely diagnosis of SCD can help in the effective management of the disease. In this article, we utilized a smartphone-based image acquisition method for capturing RBC images from the SCD patients in normoxia and hypoxia conditions. A computer algorithm is developed to differentiate RBCs from the patient’s blood before and after cell sickling. Using the developed smartphone-based technique, we obtained similar percentage of sickle cells in blood samples as analyzed by conventional method (standard microscope). The developed method of testing demonstrates the potential utility of the smartphone-based test for reducing the overall cost of screening and management for SCD, thus increasing the practicality of smartphone-based screening technique for SCD in low-resource settings. Our setup does not require any special storage requirements and is particularly useful in assessing the severity of the SCD. This is the characteristic advantage of our technique as compared to other hemoglobin-based POC diagnostic techniques.

## Introduction

Sickle cell disease (SCD) is a common worldwide genetic disorder caused by the single point mutation in the beta-globin gene [1-3]. The β-6 glutamic acid is substituted by Valine, leading to the transformation of normal hemoglobin into HbS [4]. At low levels of oxygen, the HbS polymerizes, and results in sickled shape RBCs [5, 6]. This sickling of cells makes them hard and sticky and as a result, severely affects their oxygen transport efficiency and blood circulation. Patient experiences acute vaso-occlusive pain in children as well as in adults [7, 8]. The children born in resource-limited settings are at a greater risk of SCD [9]. Centers for Disease Control (CDC) has reported about 100,000 cases of individuals with homozygous genotype from a 2008 census in the USA population mainly in African Americans [10]. SCD is the most prevalent disease in sub-Saharan Africa with the highest incidence of deaths in children under 5 years of age [11]. Approximately 700 children in Africa are born with SCD every day [2, 11]. Over half of them die due to lack of diagnosis and treatment of SCD. This disease can damage any part of the body, especially spleen [12]. Children having SCD are liable to the development of systemic infections due to loss of splenic functions. Another major organ affected in SCD is the lung [13]. Patients with SCD are at high risk of pulmonary hypertension at a very young age, which crucially increases mortality rates in children. Cerebrovascular disorders are also responsible for much morbidity and mortality with SCD [14, 15]. The most common risk factors associated with SCD are stroke and silent infarction. The likelihood of a child with SCD having a risk of stroke is 200 times greater than a healthy one with a top incidence of ischemic stroke (caused by a blood clot that blocks a blood vessel in the brain) between 2 to 5 years of age.

An early-stage detection of SCD can be quite effective in managing the disease, especially in home-based, point-of-care (POC), and other resource-limited settings [16]. Hemoglobin electrophoresis and high-performance liquid chromatography (HPLC) are gold standard methods for the detection of SCD [17]. They require heavy laboratory equipment, a continuous supply of electricity, approximately 1mL patient blood sample, and trained staff to operate and interpret the test. Other existing techniques rely heavily on the use of optical microscopes. The morphological changes in the sickle cells are observed during their oxygenated (normoxia) and deoxygenated (hypoxia) states. This use of bulky and expensive microscopes is a time-consuming process that can only be performed at a diagnostic lab. These standards are often impossible to meet in many parts of sub-Saharan Africa and other low-resource countries.

Several researchers have developed electrical impedance spectroscopy (EIS) based methods for the detection of SCD in microfluidic chips [18]. This combination of EIS and microfluidic has resulted in a reliable, accurate, and efficient method that can easily distinguish normal and sickled RBC and offers several advantages like label-free and non-invasiveness. The variations in the measured electrical impedance differential of sickle RBCs can work as a new biomarker of SCD. Another electrical impedance-based microflow cytometry technique with oxygen control seems potentially useful for SCD diagnosis [5]. Optical methods for sickle cell detection are based on the measurement of number of cells and different form factors. Electrical methods rely on the cellular dielectric properties, cell size, and difference in cell interior, Hb types and concentrations. They does not need advanced image processing. However, they do not provide information on disease severity. The equipment used for EIS (impedance analyzer) and oxygen control is relatively expensive and large in size, and not suitable for home-based and POC settings. It is vital to make sure that SCD diagnostic device must fulfill the World Health Organization’s criteria of being affordable, sensitive, specific, user-friendly, rapid and robust, equipment free, and delivered to those who need it, leading to the acronym “ASSURED” [19]. The further addition of features like real-time testing, communication of results and ease of sample collection leading to the acronym “REASSURED” can make such SCD devices even more suitable [20]. A POC SCD device developed on REASSURED Criteria can diagnose SCD in newborn babies and adults and can significantly reduce the associated pain episodes and mortality [21]. A variety of techniques such as HemeChip and µPADs have been reported in this effort. See reference [22] for a comprehensive review.

The widespread use of smartphones worldwide has opened new avenues for home and POC-based biomedical diagnostics [3]. A myriad of attachments has been developed to integrate with smartphones to enhance their imaging capabilities and observe medical conditions. A magnetic levitation-based platform was developed by researchers to detect sickled cells [3]. The developed setup eliminates the need for expensive centrifuge machines and microscopes for SCD detection. It rather utilizes magnetic levitation and a smartphone to capture the images of RBC levitating in the magnetic field using a smartphone camera. The sample is illuminated by an external LED and a lens is utilized for image enhancement purposes. Sickle cell levitation patterns are inherently different than those of normal RBC and this criterion may be used to distinguish the disease. This technique is limited to SCD detection and may be further developed to detect disease severity or sickle cell trait (SCT).

In this article, we have developed a POC and home-based portable and standalone setup for the diagnosis and treatment monitoring of SCD based on shape change in RBCs under hypoxia. It consists of a custom-designed 3D structure that can be easily attached to the smartphone camera. This setup supports an external lens to enhance the image quality along with the microchip to contain the blood sample. The sample is illuminated with an external LED and images of cells are captured using the smartphone camera. The captured images are analyzed using the computer algorithm written in MATLAB. The normal RBCs can be automatically distinguished from the sickled cells based on their morphology. The developed setup is cost-effective that significantly reduces the per-test cost and can easily be utilized in any home-based settings. It can diagnose the SCD and can also be utilized for monitoring the treatment. Using our technique, it is possible to determine the percentage of sickled blood cells and may potentially adjust the dose of medication. The whole diagnostic process can be completed within 16 minutes time with minimal user input.

## Materials and methods

### Microfluidic device design and fabrication

The microfluidic device was developed using previously reported method utilizing 1.5 mm thick Poly-(methylmethacrylate) (PMMA) sheets (McMaster-Carr, Atlanta, GA), the double-sided adhesive tape (DSA) (3M, St. Paul, MN, 20 µm thick) and microscope slides (Fisherbrand plain, pre-cleaned glass slides) [23-26]. The design for the device was developed using AutoCAD software. PMMA sheets were machined with VLS 2.30 laser cutter (VersaLaser, Scottsdale, AZ). A 20 mm × 2 mm channel was cut inside DSA sheet to hold the blood sample. The top layer provided an inlet and outlet with diameter 0.4 mm. The schematic of the microchip fabrication is shown in Figure 1 (a). The whole structure containing PMMA and DSA was affixed to a 75 mm × 25 mm glass slide to make a composite microfluidic device (Figure 1 (b)). These dimensions were purposely chosen as per the requirements of smartphone attachment. Approximately 1µl) of blood was injected into the microchannel as shown in Figure 1(c,d).

**Figure 1:**
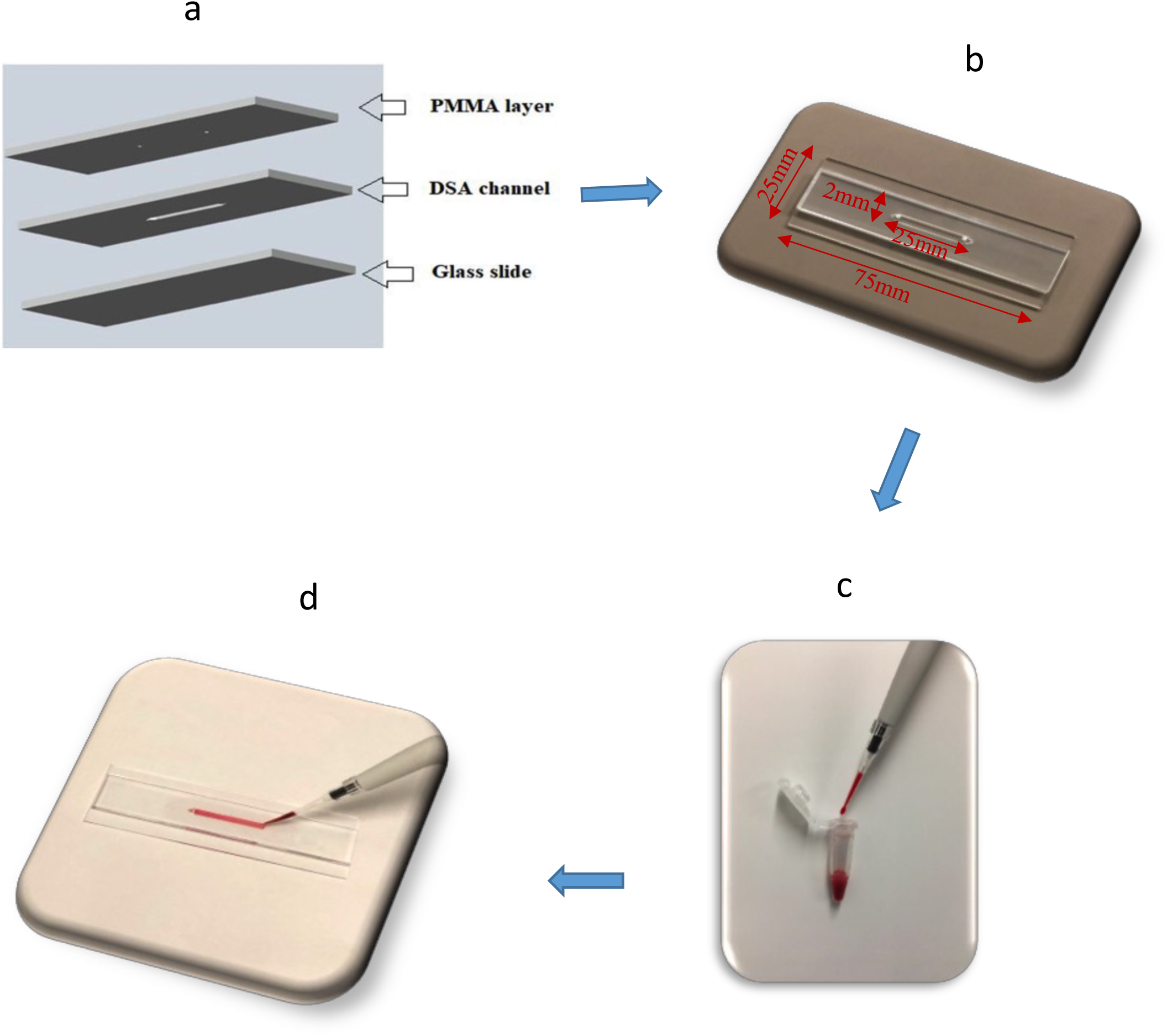
(a) Schematic illustration of microfluidic chip assembly consisting of three layers (PMMA, DSA, and glass slide) (b) Photograph of assembled microfluidic channel (length=25mm, width=2mm, height=0.02mm) (c) Blood sample used ∼1µL (d) Prepared chip containing RBCs.

### Optical attachment and smartphone-based setup

The setup was designed to operate with the rear camera of the smartphone. The optical element is composed of an aspheric lens, harvested from an internal Blu-ray drive. The lens is pasted on flexible film which cleanly sticks on the glass of any smartphone covering the camera. It was attached directly to the rear camera of the smartphone. The sample was trans-illuminated by a bright light source, powered by 2 Lithium batteries (3V each). It has the correct intensity for supplying good illumination through the samples without saturating the camera sensor. This light source is compatible with the safety standard EN62471:2008. The setup allows to recognize details of about 4.38 microns. We used the USAF 1951 slide and distinguished group 7 element 6 (Supplementary Figure S1). A stage for positioning smartphone and slide support was designed in AutoCAD. It was cut using a VLS 2.30 laser cutter (VersaLaser, Scottsdale, AZ) to support the delicate control of the focal distance and for the alignment of the light source (Figure 2). The effective focal point was established practically through trial and error. The stage was designed for a good alignment of the light source under the lens and a correct setting of the distance between lens and sample slide (1/8 inch or 3mm). The light is placed at about 20mm distance for illuminating the sickle blood sample. This distance is provided by the slide support of the smartphone stage. Figure 2 shows the top and side views for our developed smartphone-based setup detailing the smartphone positioning, slide support and alignment of light source. The dimensions of the stage were selected carefully to avoid manual focusing and errors that arise from focal distance and unstable mechanical support. The image magnification in our optical attachment was achieved using digital zoom of smartphone camera (5x with iPhone 6s). The developed smartphone system was able to image RBCs inside microchannel covering a field-of-view ∼ 0.12 mm x 0.1 mm.

**Figure 2:**
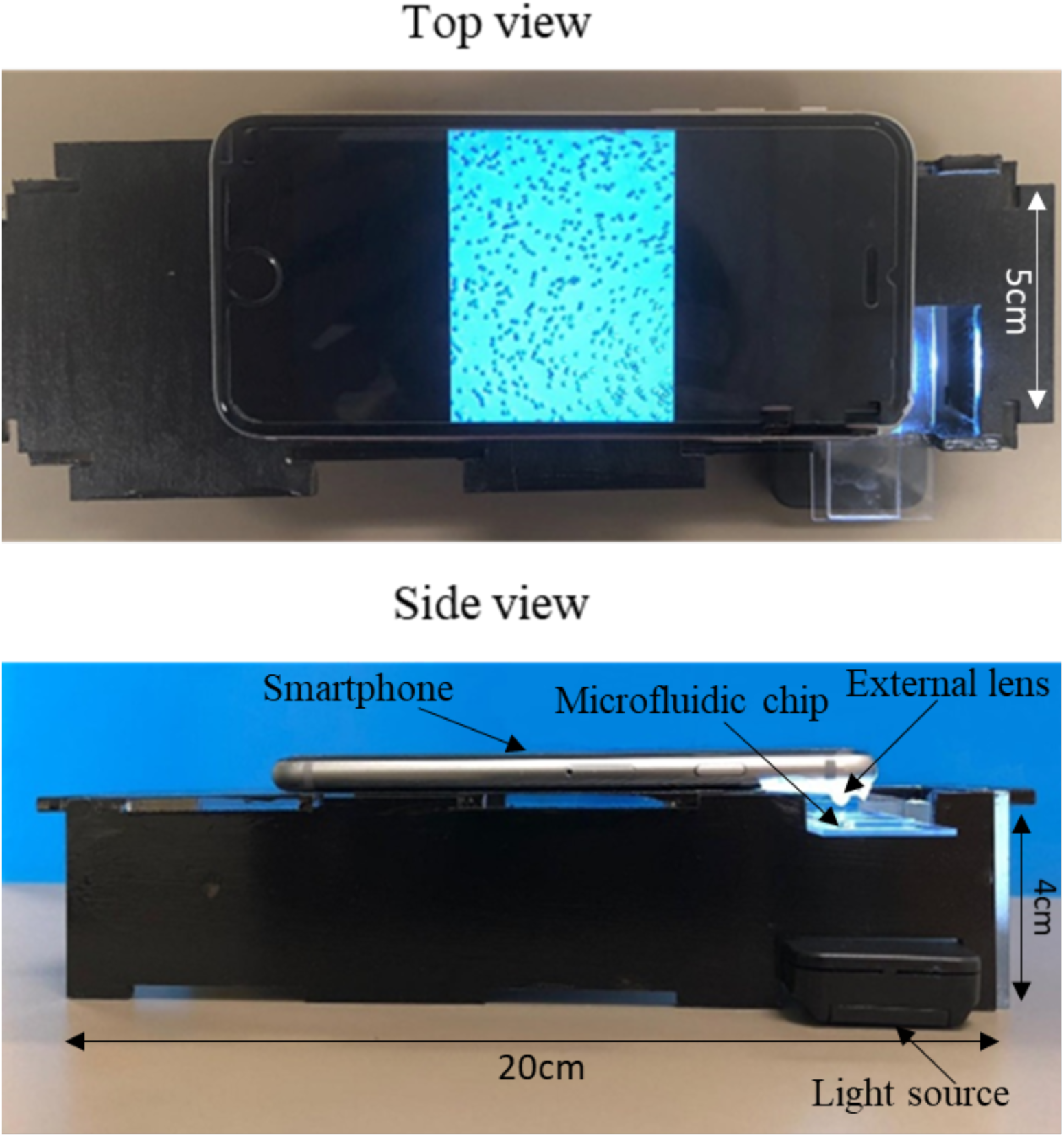
Smartphone-based platform for sickle cell blood imaging with dimensions of stage (length = 20cm, width = 5cm, height = 4cm).

### Sample preparation

De-identified whole blood from healthy donors was obtained from a local blood bank (Continental Services Group, Inc. Miami, USA). K2 anticoagulant EDTA tubes were used to collect these blood samples. De-identified sickle blood samples were obtained following Institutional Review Board (IRB) approvals from Florida Atlantic University and the University of Miami. All blood samples were diluted using phosphate-buffered saline (PBS) at a constant ratio of 1:100. We calculated the percentage of sickled cells in the blood sample obtained from two SCD patients under normal oxygen levels and hypoxia. Normally hypoxia is based on measure of; oxygen saturation in blood (SO_2_) <95%, partial pressure of oxygen (PaO_2_) <80mmHg, PH <7.35, partial pressure of carbon dioxide (PaCO_2_) >45mmHg or concentration of bicarbonate (HCO_3_) <22meq/L [27]. Less than 1 µl blood was loaded into the microfluidic device. Images were obtained using the microscope (Nikon Eclipse TE2000-S) & smartphone setup and labeled as “Before treatment.” Sodium metabisulfite (2% solution) was used to deoxygenate the RBCs (0.2 g of sodium metabisulfite in 10 ml of nano-pure water) [4, 28]. Sodium metabisulfite is a reducing agent which promotes sickling [29, 30]. The diluted blood from the normal and sickle blood samples was mixed separately into the prepared sodium metabisulfite controlling the dilution factor as 1:100. This sample was incubated at room temperature for 15 minutes and injected into another microfluidic chamber. Sodium metabisulfite reduces the oxygen tension inducing the typical sickle-shape to RBCs. Images were taken for these samples with deoxygenated hemoglobin inside RBCs using the microscope and the developed smartphone setup for comparison.

### Image processing and algorithm development

Computer-based image processing techniques are developed for counting different types of erythrocytes according to their variable morphology [31]. In this work, MATLAB R2019a is used for image analysis and quantification of sickled cells with a single click using an adaptive thresholding algorithm. Visual representation of our developed computer-based program can be seen in Figure 3. The MATLAB codes can be easily ported to the iOS/Android as well. The algorithm developed here can assess the concentration of sickled cells under hypoxia and normoxia conditions. The image processing time to calculate the percentage of sickled cells in a sample using the smartphone system is <15 seconds.

**Figure 3:**
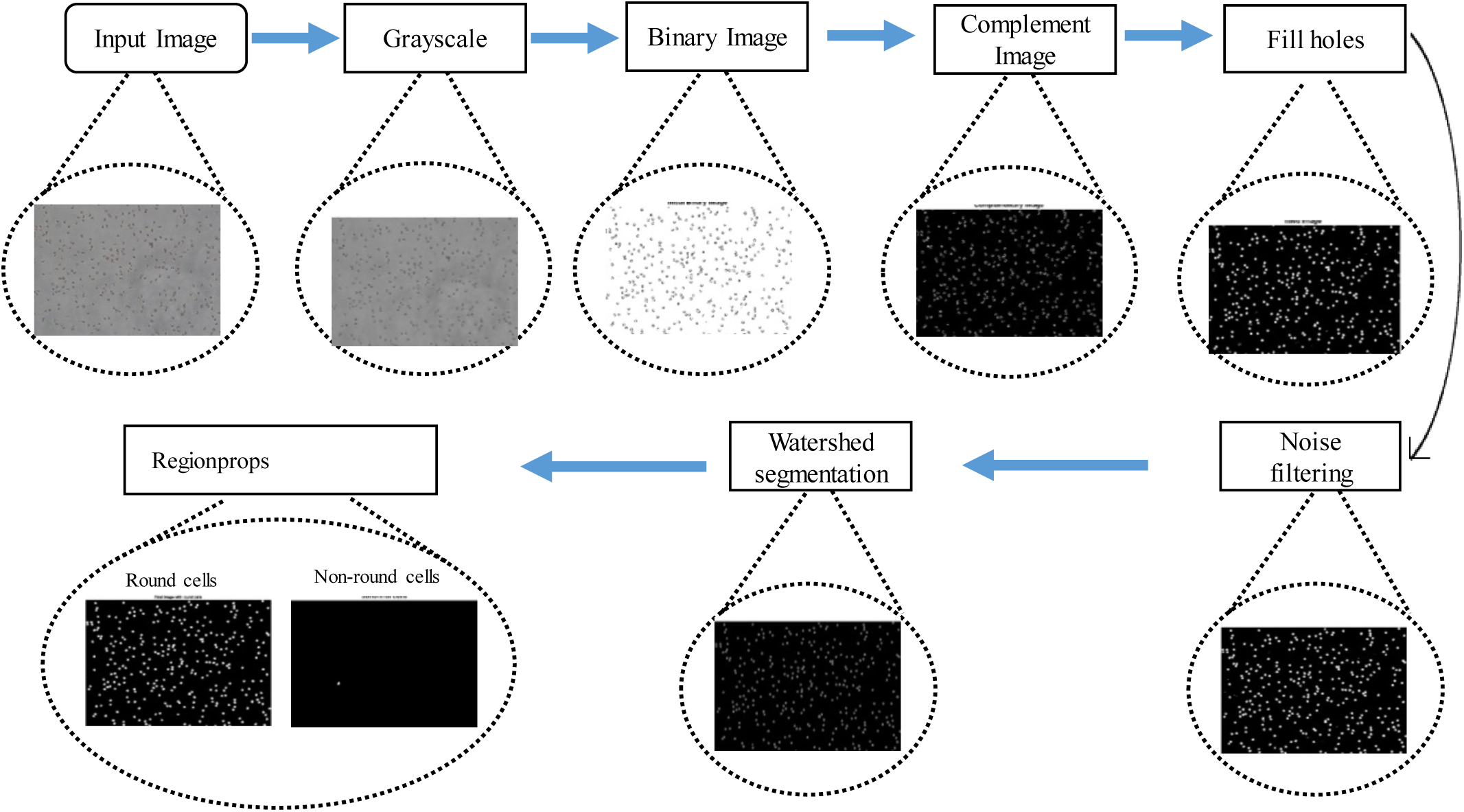
Diagram detailing the MATLAB architecture for diseased cells counting using regionprops function (microscope image).

The algorithm developed in this testing is described below:

Step 1. The human RBCs image captured either with a microscope or smartphone is fed to the program as input and is first converted to gray and then binary image using an adaptive thresholding that separates the foreground from the background with nonuniform illumination.

Step 2. The binary image is complemented, and the small holes inside the objects are filled up using imfill operator to increase the accuracy of the further calculation.

Step 3. For the ease of further processing, cells on the borders are removed using imclearborders.

Step 4. The watershed transform is used to segment adjacent RBCs into separate bodies. The watershed transforms “watershed ridge lines” in an image by treating it as a surface where light pixels represent high elevations, and dark pixels represent low elevations.

Step 5. Convex hull image is generated from the binary image using bwcovhull.

Step 6. Small unnecessary spots are removed from the image using bwareaopen operation. It eliminates all the objects in the diagram containing smaller number than the number of pixels mentioned in the threshold level.

Step 7. Connected components were labeled in a 2-D binary image using bwlabel.

Step 8. Area and perimeter of each of the components is calculated using regionprops on the objects.

Step 9. Discarding the smaller background objects, only the larger RBCs are considered and pre-processed for going through the rest of the steps.

Step 10. Metric [i.e. (perimeter^2^/4*π*area)] is calculated for each object. A circle, with radius r has an area of π*r^2^ and a perimeter of 2*π*r. A common place resolution independent measure of roundness is calculated by:

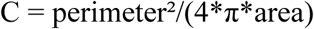

where C is the measure of circularity. If the value of C is equal to one, its circle, and if not, then it has other shapes [32, 33].

Step 11. By this metric value, any deviation or change in the shape of RBC is detected, which gives the number of sickled cells in the sample.

## Results

The smartphone setup used for screening sickled cells is illustrated in Figure 2. The whole platform comprised of an aspheric lens, printed stage, and a light source to capture the images of RBCs inside microfluidic channel. The stage was designed to stabilize the smartphone upright and the sample was input into microfluidic chamber through an inlet. The smartphone camera captured images of the microfluidic chip through the aspheric lens that is pasted on the smartphone camera lens to focus the RBCs. An LED under the channel is used to enhance imaging. It can be easily modified for other smartphones. Blood samples were diluted using PBS and images were captured under microscope and smartphone setup to make a comparison between the results. 2% sodium metabisulfite solution (reducing agent) was prepared and mixed with the samples in separate eppendorfs and imaged under microscope and smartphone likewise.

To evaluate the optical attachment setup and MATLAB-based program, first we performed the experiment using normal blood samples from healthy donors. Validation image results were obtained with two healthy blood samples using 3 different areas for field-of-view each covering 0.12 mm x 0.1 mm (∼0.036 mm^2^ total area) for all samples. Images were captured with our mobile setup and laboratory-grade standard microscope for direct comparison. MATLAB algorithm performed the necessary image processing. Our system automatically executed cell classification and calculated the number of total cells, extracted sickled cells, and normal cells. For the normal blood sample 1, a very small average number of healthy blood cells (0.2%±0.08) were calculated as abnormal/sickled cells before treatment using the microscope images and 0.26%±0.09 cells were classified as abnormal cells after treatment (false-positive results). Likewise, for cellphone images, 0.86%±0.12 and 3.6%±1.3 healthy RBCs were misclassified as sickled cells pre and post-treatment, respectively. Similarly, for the second normal blood sample, 0.3%±0.14 cells were classified as sickled before treatment using microscopic images, and 0.45%±0.17 cells were detected as sickled after treatment. In images taken with smartphone 1.65%±0.77 and 2.9%±1.22 cells were counted as sickled before and after treatment, respectively. Our developed system calculated an insignificant number of sickled cells for known healthy blood samples and presented comparable results for both types of images. The percentage of sickled cells calculated by our platform for each one of these normal blood samples is listed in Table 1, and an example of image transformation through different digital processing steps is also shown in Figure 4.

**Table 1:** Result set for the percentage of sickled blood cells.

| Serial # | Percentage of sickled cells |  |  |  |  |  |  |  |
| --- | --- | --- | --- | --- | --- | --- | --- | --- |
|  | Normal blood sample 1 |  | Normal blood sample 2 |  | Patient 1 |  | Patient 2 |  |
|  | Smartphone | Microscope | Smartphone | Microscope | Smartphone | Microscope | Smartphone | Microscope |
|  | Image | Image | Image | Image | Image | Image | Image | Image |
| Before treatment |  |  |  |  |  |  |  |  |
| Image 1 | 1% | 0.3% | 1.7% | 0.23% | 6.9% | 6% | 2.4% | 2.8% |
| Image 2 | 0.9% | 0.1% | 0.86% | 0.3% | 5.4% | 5.6% | 5.3% | 5.5% |
| Image 3 | 0.7% | 0.2% | 2.4% | 0.5% | 6.7% | 6.8% | 3.5% | 2% |
| Average | 0.86%±0.12 | 0.2%±0.08 | 1.65%±0.77 | 0.3%±0.14 | 6.3%±0.66 | 6.14%±0.49 | 3.7%±1.19 | 3.4%±1.49 |
| After treatment |  |  |  |  |  |  |  |  |
| Image 1 | 2.9% | 0.2% | 3.5% | 0.26% | 37% | 35.6% | 45.4% | 40.5% |
| Image 2 | 5.7% | 0.4% | 3.7% | 0.5% | 38.1% | 32.6% | 41% | 40.7% |
| Image 3 | 2.9% | 0.2% | 1.5% | 0.6% | 34% | 36.13% | 44% | 40.4% |
| Average | 3.6%±1.3 | 0.26%±0.09 | 2.9%±1.22 | 0.45%±0.17 | 36.3%±1.7 | 32.7%±1.5 | 43.7%±1.4 | 40.5%±0.12 |

**Figure 4:**
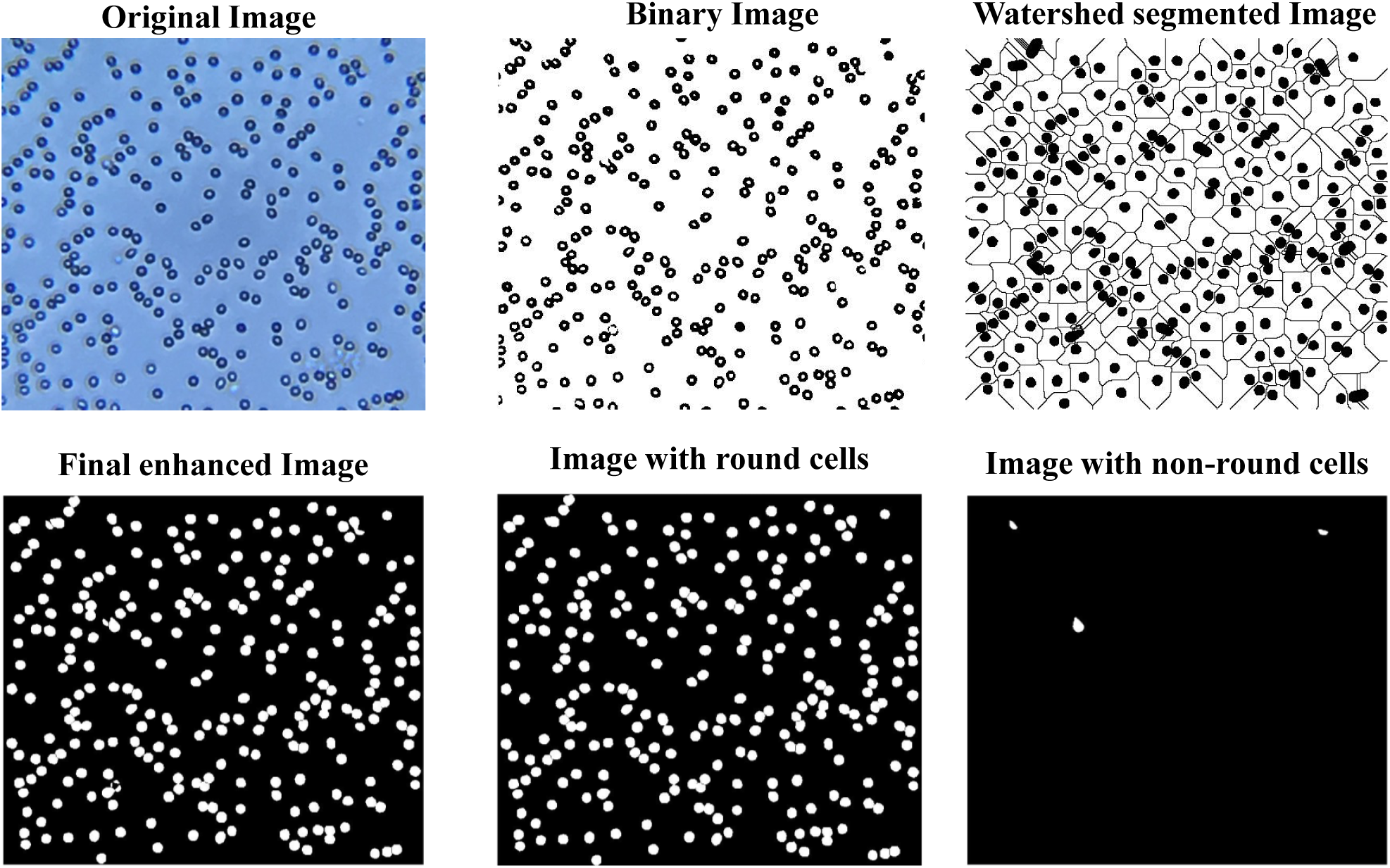
Normal blood cells image captured using smartphone-based setup before treatment.

Following the validation of our developed framework, we performed the same testing technique using blood samples from two SCD patients (patient 1 & patient 2). Three different patches inside the channel were imaged covering a total area of approximately 0.036mm^2^. The system classified and counted RBCs using the clinical blood samples before deoxygenation and presented a different set of results. An average of 6.1% and 6.3% sickled cells for patient 1 blood sample and an average of 3.4% and 3.7% for patient 2 blood sample indicated the deformation of cells in a very low proportion under normal oxygen levels. A significant number of sickled cells were obtained using post-treatment clinical samples from both the patient 1 & 2. Computing the values of percentages of sickled cells, system gave us an average of 32.7% ± 1.5 and 36.3% ± 1.7 for two types of images (patient 1). The percentage values for patient 2 blood sample were also consistent for microscope and smartphone images (40.5% ± 0.12 and 43.7% ± 1.4 respectively). Using our platform, patient 1 and patient 2 blood samples were classified as SCD positive. Supplementary Figure S2 represents examples of sickle blood sample images using a smartphone-based setup and microscope for the same sample. The percentage of sickled cells after deoxygenation is associated with predictable physiologic and clinical parameters and can be used as a measure of SCD severity and patient’s disease outcome [34]. Our developed system has the potential to be used as a screening tool for SCD and other blood cell disorders in resource-limited settings.

## Discussion

Here, we present the development and evaluation of a smartphone-based optical setup for the detection and quantification of sickled cells using disposable microfluidic chip containing a small volume of blood ∼1µL. The main advantages of using microchip are the ease of utilization, the requirement of only fingerprick blood volume and high uniformity of blood cells in microfluidic device. Blood smears are frequently utilized in hematological analysis where the uniformity of the cells may be affected by the expertise of the operator [35]. We imaged the evenly distributed RBCs in the microfluidic chip with our developed smartphone-based setup and compared the results with the optical microscope image results. We have used the blood from normal & SCD patients and captured the images under normal and hypoxic conditions using our smartphone-based setup and microscope. A program written in MATLAB counted number of cells for each kind and classified them based on their shapes to count percentages of sickled cells. The blood samples containing more than a significant number of sickled cells were categorized as being SCD positive. If the average percentage number of sickled cells was insignificant, the sample was classified as normal. Our results are comparable with the results obtained using microscope for the same sample. For example, for patient 1, we obtained a percentage of 6.3% sickled cells under normal conditions and 36.3% sickled cells under hypoxic conditions. Using optical microscope images 6.1% sickled cells before treatment and 32.7% sickled cells after treatment were calculated which are consistent with the results obtained using our developed method.

Previous studies have shown a close relationship between sickle red blood cells (SS-RBCs) morphology and hemoglobin polymers aligned inside the cells [36]. The maximum sickled fraction indicated a strong positive correlation with the HbS percentage level during the long-term deoxygenated state [37]. It is proved that there is a clear inverse correlation between the percentage of sickled erythrocytes under hypoxia and fetal hemoglobin (HbF) levels [34]. Percentage of sickled RBCs after deoxygenation are correlated negatively with pH and positively with the presence of long fibers inside the erythrocytes. Environmental factors, such as pH, and patient related factors, such HbF levels are well known to influence the rate of HbS polymer formation [38, 39]. HbS polymer fraction is associated directly with disease severity [40]. Decrease in pH may lead to painful crisis in SCD patients and treatment with alkali administration may be beneficial [41]. Hydroxyurea has been the only Food and Drug Administration (FDA) approved drug to treat SCD in adults from previous two years [42, 43]. It reduces the frequency of painful episodes and enhances the amount of HbF and hemoglobin. Recently, FDA approved Adakveo® (crizanlizumab-tmca) medicine to reduce the frequency of painful crises in adult and pediatric patients with SCD [44-46]. This is an important advancement for people living with this disease. Adakvo binds to a cell adhesion protein called P-selectin that plays a vital role in multicellular interactions that can activate vaso-occlusion [8, 47].There is no useful drug reported to prevent or reverse the polymerization of HbS [48].

This POC technology can potentially be applied to diagnose and improve SCD management in resource-limited settings [22]. Most of the laboratory-based technologies are expensive and limited to testing centers [49]. The developed technology has the potential to reduce the costs significantly for SCD testing and care. The smartphone-based system presented here requires a mass-producible and economical microfluidic device with an optical smartphone attachment with overall cost less than $10. The total material cost to fabricate the microfluidic chip was approximately <$1, which includes $0.1 for PMAA, ∼10 cents for the double-sided adhesive (DSA), $0.10 for glass slide. The material cost to fabricate the smartphone attachment was <$7, with ∼$1 for the 3D printed smartphone accessory, $5.91 for an LED, $1 for the lenses, and $2 for the battery. Sodium metabisulfite for a single test cost is ∼$0.01. This assay is user-friendly and does not require any trained personnel and can potentially be self-performed. A drop of blood can be mixed using disposable transfer pipette with PBS. A 15-minute waiting period is required, after mixing blood with sodium metabisulfite. Loading the sample into microfluidic channel, capturing the image with smartphone, and one click analysis can be performed in less than a minute. The sodium metabisulfite can potentially be dried inside microfluidic device for self-testing purpose if needed.

Early-stage diagnosis and screening of SCD can significantly improve the management of disease, particularly in resource-constrained areas. Critical complications, including stroke, organ damage, blindness, leg ulcers, and others can be prevented if proper management of the disease is done and mortality rate can be drastically reduced. Our system can rapidly quantify the sickled RBCs. Based on its properties, each cell is examined and classified as normal/sickled. This system has potential for screening of SCD and its management.

## Conclusion

We have demonstrated a simple, rapid, and cost-effective smartphone-based SCD detection method. This developed platform utilizes an external lens that can be easily attached to the smartphone camera to record images of various blood samples inside a microchip. The captured images are rapidly processed using a MATLAB program and the total number of sickled cells is automatically counted. To evaluate the performance of our setup, we used normal blood sample as well as the SCD patients’ samples. Our developed image processing algorithm accurately quantified sickled cells in deoxygenated blood. The smartphone-based quantification results were compared and validated with the optical microscope data. With its simple sample preparation combined with small volume, our developed setup is well-suited for resource-limited settings providing remote diagnosis opportunities.

## Acknowledgments

We acknowledge research support from NIH R15AI127214, R56AI138659, Institute for Sensing and Embedded Networking Systems Engineering (I-SENSE) Research Initiative Award, FAU Faculty Mentoring Award, Humanity in Science Award, and a start-up research support from College of Engineering and Computer Science, Florida Atlantic University, Boca Raton, FL.

