## Supplemental Information for "Smartphone-based Sickle Cell Disease Detection and Monitoring for Point-of-Care Settings"

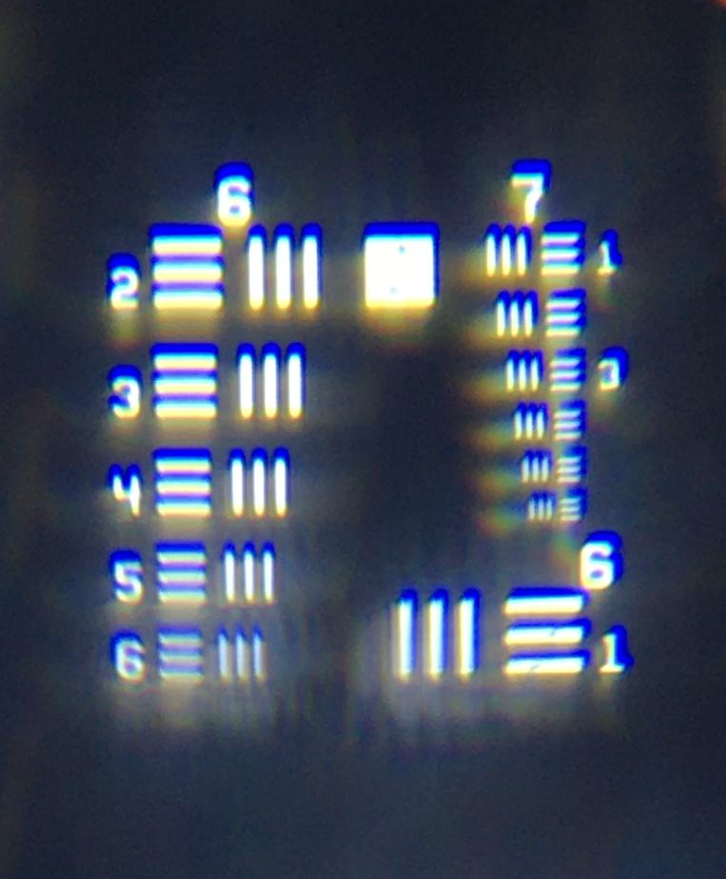

**Supplementary Figure 1.** USAF 1951 slide imaged using our developed setup.

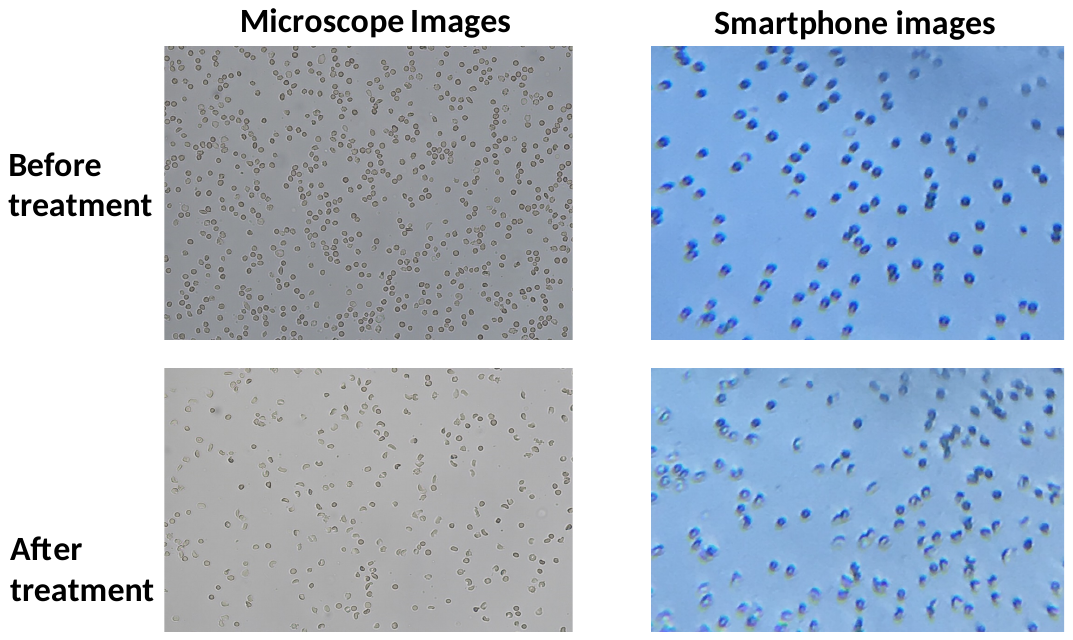

**Supplementary Figure 2.** SCD blood images used for direct comparison between smartphone and microscope image analysis results.
